# Is this a fake QTL? A short note on analysis of multiple QTL peaks on the same chromosome

**DOI:** 10.1101/270736

**Authors:** Mathias Lorieux

## Abstract

A simple method is proposed to detect fake QTL provoked by spurious linkage with a true QTL. The method is based on the calculation of the expected value of the teststatistic (F-test and LOD score are considered) at the putatively fake QTL position.

## The problem

In quantitative trait locus (QTL) mapping, it is frequent to observe several peaks in the plot of the test statistic (LOD score, F-test, etc.) against the linkage group position - the statistic decays sufficiently between local peaks to allow declaring each peak as a different QTL. In this case, one can legitimately ask if some of the peaks are actually spurious QTLs generated by, for example, a departure of the data from its expected structure model. Distant genomic regions on the same chromosome that show unexpected linkage strength are an example of such departures. Methods like Multiple Interval Mapping (MIM) (Jansen 1993; Kao *et al.* 1999), Composite Interval Mapping (CIM) ( Zeng 1994) or Inclusive Composite Interval Mapping (ICIM) (Li *et al.* 2007) are designed to co-analyze several QTL peaks, however they might not be adequate for such situations (I’d like to leave this point open for discussion).

## Method

For simplicity, let’s consider the case of two QTL peaks found on a linkage group for the same trait. Suppose an F-test statistic has been obtained from the regression of phenotypes onto genotypes in a segregating population, and then plotted along a chromosome at every marker position. Suppose also that the plot shows a main peak that reveals the presence of a QTL, but also seems to reveal a second, lower QTL peak, at some distance from the main peak. One needs to decide if the secondary QTL peak is actually due to a real QTL effect, or if it is simply the result of linkage with the main QTL. Normally, if there was only one *real* QTL on a chromosome, one would expect to observe a propagation effect from the main QTL: the F-test value should decrease continuously with the genetic distance (i.e., the recombination fraction) from the true QTL position. Moreover, no other QTL local peak should show up, apart from small stochastic variations due, for example, to missing or erroneous data at some marker positions. But this is only true under the assumption that the recombination odds are following their expected values.

Sometimes, abnormally tight linkage between markers located in distant chromosomal regions can be observed. This can be due to different causes - for example, epistatic interaction of sterility genes. One can consider this situation as a special case of inadequacy between the data structure and the data model used in the QTL analysis, where the data model says that the pairwise recombination fraction between any pair of markers, and the sum of adjacent recombination fractions (SARF) of all the intervals that separate the two markers are equivalent, while in the real data they are *not*. In such cases, one could question if the secondary QTL peak is primarily explained by its spurious linkage with the main QTL peak-region. This hypothesis could be tested by a careful inspection of the *expected* value of the F-test at a given marker position, *E*(*F*_*M*_).

A way to calculate *E*(*F*_*M*_) is as follows. Consider a QTL *Q* and a marker *M* linked to *Q*. Let *F*_*Q*_ be the observed value for the F-test at the *Q* position, and 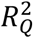 be the fraction of the trait variation explained by *Q*. Let also *r* be the recombination fraction between *Q* and *M*, *n* be the population size and *p* be the number of genotypic classes observed in the marker. Then, we know that the expected value of *R*^*2*^ at any marker *M* is

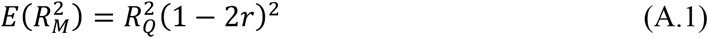

Since we know from linear regression theory that

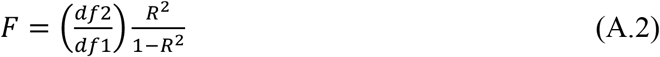

which leads to

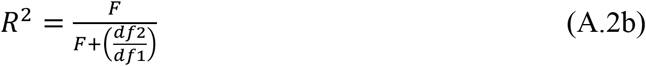

where *df* 2 = *n − p* and *df* 1 = *p* − 1 are the degrees of freedom used to calculate *F*, we can plug (A.2b) in (A.1) to derive the expected value of the F-test at marker *M*

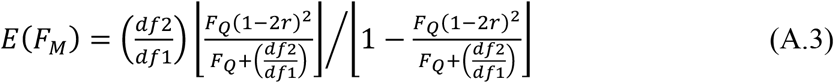

which slightly simplifies to

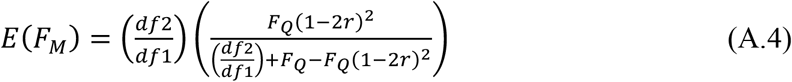

In backcross or double haploid populations, there are only two genotypic classes, so *df*2 = *n* − 2 and *df*1 = 1, and

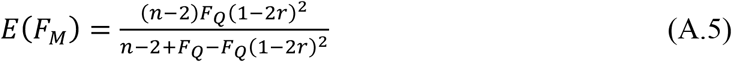

In a recombinant inbred line (RIL) population derived by single-seed descent, *r* is replaced by 2*r* /(1 + 2*r*) in (A.5).

In F_2_ populations, three genotypic classes are observed, so *df*2 = *n* − 3 and *df*1 = 2, and

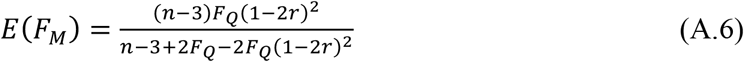

To test if a second, smaller QTL is actually a fake one, we compare two different values for *E*(*F*_*M*_), *E*_*p*_(*F*_*M*_) and *E*_*d*_(*F*_*M*_), with the observed F-test values (by plotting all three values against the marker positions on the chromosome, expressed in centimorgans or cM). *E*_*p*_(*F*_*M*_) and *E*_*d*_(*F*_*M*_) are calculated using two different estimates for *r*, respectively: the first one, 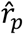, is the observed *pairwise* recombination fraction between marker *M* and the marker that has the highest F-test, *M*_*Q*_, and the second one, 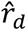, is derived from the sum *d* of the sizes in cM of the adjacent intervals defined by the markers that lay between *M* and *M*_*Q*_. 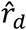 is then obtained from *d* by inverting the mapping function - e.g., Haldane’s or Kosambi’s function - that was used to calculate the genetic map. In case of spurious linkage, 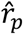 can be significantly lower than 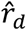, which is likely to be closer to the true recombination fraction that *would* be observed in a population that follows the expected values. Note that the markers are assumed to be densely distributed within the *M* - *M*_*Q*_ interval, so we avoid the “ghost QTL” effect (see Haley & Knott 1992; Martínez & Curnow 1992); this is important because we need the main QTL to be precisely located.

The difference between the *E*_*p*_(*F*_*M*_) and *E*_*d*_(*F*_*M*_) values reflects the effect of the discrepancy between 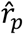 and 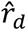 on *E*(*F*_*M*_). Consequently, if the second, smaller QTL is due to spurious linkage with the main QTL, we expect the curves for the observed F-test and its expectation *E*_*p*_(*F*_*M*_) to have similar shapes and close values, and *E*_*d*_(*F*_*M*_) to show a regular decrease from the QTL peak and an obvious departure from *E*_*p*_(*F*_*M*_). This departure could be tested by calculating the probability log-ratio

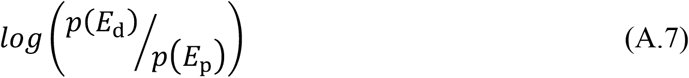

The expected LOD score at QTL position *M*, *E*(*Z*_*M*_), can be derived using a similar approach. We can use the generalized *R*^*2*^ expression, proposed by Nagelkerke (1991), to derive the relationship between *R*^*2*^ and *Z*

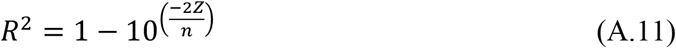

which leads to the following expression for *E*(*Z*_*M*_)

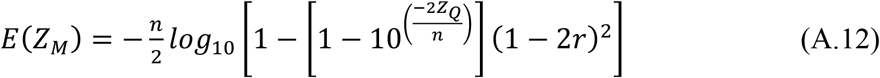

where *Z*_*Q*_ is the LOD score statistic at QTL position *Q*.

Then we can calculate *E*(*Z*_*M*_) for 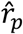 and 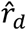 just as we did for *E*_*p*_(*F*_*M*_) and *E*_*d*_(*F*_*M*_), and compare the curves to the observed LOD score in order to conclude on the presence of a fake QTL. Note, (A.12) is valid for any population type.

Figure 1 illustrates an application of the method to a QTL study of resistance to the parasitic plant *Striga hermonthica* in rice.

**Figure 1.**
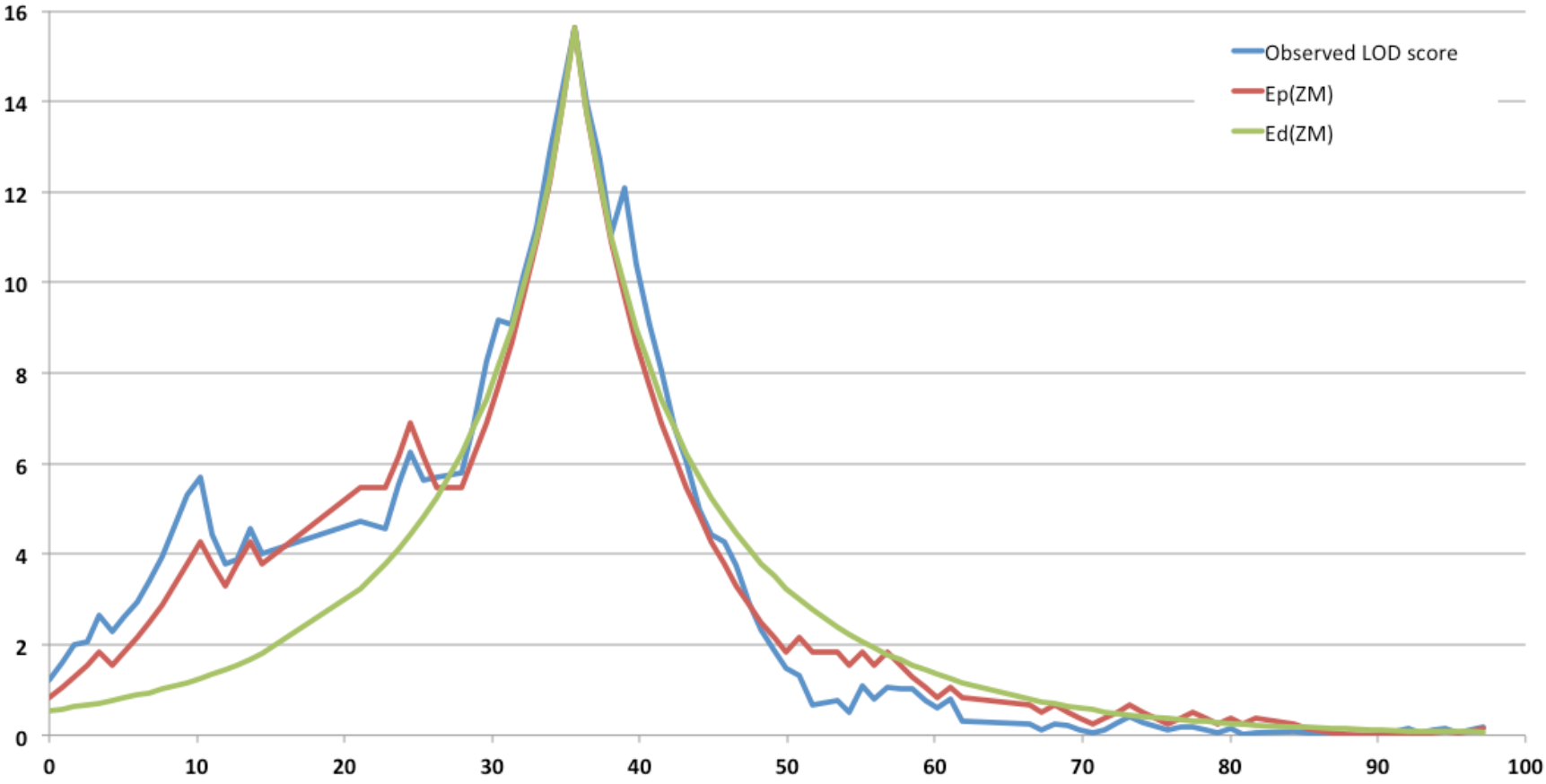
Observed and expected LOD score, for resistance to *Striga hermonthica* along a rice chromosome, in a RIL population. A secondary peak is observed at ^~^10 cM. Observe the resemblance of shape between the observed LOD score and its expected value *E*_*p*_(*Z*_*M*_) calculated using pairwise recombination fractions between the QTL peak and every marker position. Also observe the difference between *E*_*p*_(*Z*_*M*_) and *E*_*d*_(*Z*_*M*_) at ^~^10 cM. We thus declare the secondary peak as a fake QTL.

## Conclusion

The proposed method allows detecting a fake QTL induced by spurious linkage with a true QTL. It can be used as a complement to other methods like CIM, ICIM or MIM as an additional test for QTL presence. We haven’t considered the case of a fake QTL provoked by spurious linkage with a true QTL located on a different chromosome, however we don’t see a reason why the method couldn’t be applicable to this case.

***Additional note.*** The sometimes used, simpler formula *E*_*b*_(*F*_*M*_) = *F*_*Q*_(1 − 2*r*)^2^, although biased, remains a fair approximation of *E*(*F*_*M*_) for large populations since the ratio between the two estimates is, for populations with two genotypic classes

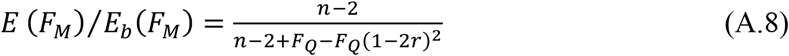

and *n* is usually much larger than *F*_*Q*_. Nevertheless, the quality of the approximation decreases as the contribution of the QTL to the trait increases.

However the bias of *E*_*b*_(*F*_*M*_) can be quite serious for smaller *n* values and highly significant QTL, especially when *r* is large. This bias is, after simplification

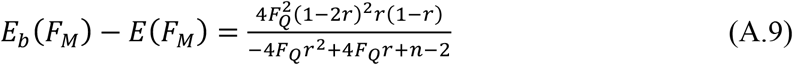

And the relative bias is

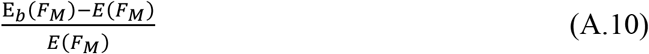

For example, in a backcross population of *n* = 200 individuals, if *r* = 0.45 and *F*_*Q*_ = 40, the relative bias is 20%, and it increases to an unacceptable ~ 40% if *n* = 100.

## Acknowledgements

Many thanks to Julie Scholes, James Bradley and Jon Slate from the University of Sheffield (UK) for their valuable comments and edits on the manuscript.

